# Circulating microRNAs as noninvasive biomarkers for canine Cushing’s syndrome

**DOI:** 10.1101/2021.08.17.456598

**Authors:** Karin Sanders, Anouk Veldhuizen, Hans S. Kooistra, Adri Slob, Elpetra P.M. Timmermans-Sprang, Frank M. Riemers, Sylvie Daminet, Federico Fracassi, Sebastiaan A. van Nimwegen, Björn P. Meij, Sara Galac

## Abstract

Canine Cushing’s syndrome (hypercortisolism) can be caused by a pituitary tumor (pituitary-dependent hypercortisolism; PDH) or a cortisol-secreting adrenocortical tumor (csACT). For both cases, noninvasive biomarkers that could pre-operatively predict the risk of recurrence after surgery would greatly impact clinical decision making. The aim of this study was to determine whether circulating microRNAs (miRNAs) can be used as noninvasive biomarkers for canine Cushing’s syndrome.

After a pilot study with 40 miRNAs in blood samples of healthy dogs (*n* = 3), dogs with PDH (*n* = 3) and dogs with a csACT (*n* = 4), we selected a total of 20 miRNAs for the definitive study. In the definitive study, these 20 miRNAs were analyzed in blood samples of healthy dogs (*n* = 6), dogs with PDH (*n* = 19, pre-and post-operative samples) and dogs with a csACT (*n* = 26, pre-operative samples).

In dogs with PDH, six miRNAs (miR-122-5p, miR-126-5p, miR-141-3p, miR-222-3p, miR-375-3p and miR-483-3p) were differentially expressed compared to healthy dogs. Of one miRNA, miR-122-5p, the expression levels did not overlap between healthy dogs and dogs with PDH (*p* = 2.9×10^−4^), significantly decreased after hypophysectomy (*p* = 0.013), and were significantly higher (*p* = 0.017) in dogs with recurrence (*n* = 3) than in dogs without recurrence for at least one year after hypophysectomy (*n* = 7). In dogs with csACTs, two miRNAs (miR-483-3p and miR-223-3p) were differentially expressed compared to healthy dogs. Additionally, miR-141-3p was expressed significantly lower (*p* = 0.009) in dogs with csACTs that had a histopathological Utrecht score of ≥ 11 compared to those with a score of < 11.

These results indicate that circulating miRNAs have the potential to be noninvasive biomarkers in dogs with Cushing’s syndrome that may contribute to clinical decision-making.

## 1 Introduction

Spontaneous Cushing’s syndrome, or hypercortisolism, is one of the most commonly diagnosed endocrinopathies in dogs (1). It is caused by an ACTH-secreting pituitary tumor (pituitary-dependent hypercortisolism; PDH) in ∼80-85% of cases, and by a cortisol-secreting adrenocortical tumor (csACT) in ∼15-20% of cases (1). Both PDH and csACT can be treated by surgically removing the causative tumor. Because surgery is not without risks and not suitable for every patient, dogs with Cushing’s syndrome are often treated with the steroidogenesis inhibitor trilostane (2). Although trilostane can effectively reduce the clinical signs associated with hypercortisolism, it does not inhibit tumor growth (3).

Pituitary tumors are usually classified as adenomas, but can nonetheless compress or invade surrounding tissues (4). After hypophysectomy, recurrence of hypercortisolism occurs in ∼23-27% of cases (5,6), for which the most important predictive marker is the pituitary height to brain area value (P/B value) (6). To determine the P/B value, diagnostic imaging with CT or MRI is necessary. The mRNA expression of pituitary tumor transforming gene 1 (*PTTG1)* was previously reported to be associated with disease-free interval after surgery, but can only be assessed post-operatively (7).

A csACT can be classified as an adrenocortical adenoma (ACA) or an adrenocortical carcinoma (ACC) (8), but making this distinction can be difficult (9). After adrenalectomy, recurrence of hypercortisolism occurs in ∼30-38% of cases (9,10), but predicting recurrence remains challenging. We have recently developed a novel histopathological scoring system, the Utrecht score, to predict the prognosis of dogs after adrenalectomy (9). In addition, we have identified three genes of which the mRNA expression in csACTs was associated with survival after adrenalectomy (11). However, these markers can only be assessed when the csACT has already been surgically removed. The only known pre-operative marker that is associated with survival after adrenalectomy is the tumor diameter, but has low predictive value (9,12).

Having noninvasive biomarkers that can pre-operatively predict the risk of recurrence after surgery would greatly impact clinical decision making. Potential candidates for such noninvasive biomarkers are circulating microRNAs (miRNAs; miRs). MiRNAs are single-stranded, non-coding RNAs of ∼20-24 nucleotides in size (13). They function as antisense RNAs which regulate their target genes post-transcriptionally, and can influence cellular differentiation, proliferation and apoptosis (13,14). MiRNA expression patterns can be altered in multiple diseases, including cancer. MiRNAs that are expressed higher in cancer are regarded as oncomiRs, while miRNAs that are expressed lower in cancer are regarded as tumor suppressor miRNAs (13). Most miRNAs are expressed intracellularly, but numerous miRNAs can also be found in biological fluids such as blood, urine, saliva, and cerebrospinal fluid. These miRNAs are referred to as circulating miRNAs (15). MiRNAs that are detectable in plasma or serum samples are bound with lipid proteins or encapsulated in extracellular vesicles and are therefore resistant to RNase digestions (13). Consequently, their expression levels are remarkably stable, which makes them ideal candidates as noninvasive biomarkers for various diseases (13).

The aim of this study was to determine whether circulating miRNAs can be used as noninvasive biomarkers for canine Cushing’s syndrome. We aimed to identify miRNAs associated with the presence or absence of PDH or a csACT, the size of the pituitary tumor and whether PDH recurred after hypophysectomy, and the histopathological assessment of a csACT.

## 2 Materials & Methods

### 2.1 Pilot study

#### 2.1.1 Animals and samples

For the pilot study, we included blood samples of healthy dogs (*n* = 3), dogs with csACTs (*n* = 4), and dogs with PDH (*n* = 3). Written informed consent was obtained from the owners for the participation of their animals in this study. Serum samples were available from the healthy dogs and dogs with csACTs, while EDTA plasma samples were available from the dogs with PDH. After sample collection, serum or EDTA plasma was obtained by centrifuging whole blood for 5 min at 4000 rpm. The serum and EDTA plasma samples were stored at −70°C until use.

The suspicion of hypercortisolism was based on the dogs’ medical history and findings during physical examination. The diagnosis of PDH was made by demonstration of suppressible hypercortisolism with endocrine testing (low-dose dexamethasone suppression test or urinary corticoid to creatinine ratios (UCCR) combined with high-dose dexamethasone suppression test), or non-suppressible hypercortisolism combined with findings of symmetric bilateral adrenal enlargement (indicating bilateral hyperplasia) and the absence of unilateral adrenal enlargement (indicating an ACT) on ultrasonography or CT, together with findings of pituitary enlargement on CT or MRI. The diagnosis of a csACT was made by non-suppressible hypercortisolism demonstrated with endocrine testing, combined with the presence of an ACT found on ultrasonography or CT. All pituitary tumors were classified as adenomas based on histopathological assessment and absence of detectable metastases (4). Histopathological assessment of the ACTs confirmed their adrenocortical origin, and the histopathological Utrecht score (Ki67 proliferation index (PI) + 4 if ≥ 33% of cells have clear/vacuolated cytoplasm + 3 if necrosis is present) was used to predict the risk of recurrence (9). The dogs in the “healthy” group were regarded as healthy based on the absence of clinical signs, no abnormalities in complete blood count and blood biochemistry, and UCCRs within reference values.

#### 2.1.2 MiRNA isolation and cDNA synthesis

MiRNA was isolated using the miRNeasy Serum/Plasma Kit (Qiagen, Venlo, the Netherlands) according to the manufacturer’s instructions. The serum and EDTA plasma samples were thawed on ice. As sample input 150 µL was used, and volumes of the other components were adjusted accordingly. The miRNeasy serum/plasma kit was combined with the RNA Spike-In Kit, For RT (Qiagen) for miRNA isolation quality control. The mixture of spike-ins UniSp2, UniSp4 and UniSp5 was prepared according to the manufacturer’s instructions, and 1 µL of the spike-in mixture was added per sample. After isolation, the miRNA eluates were stored at −20°C until use. For subsequent cDNA synthesis, miRNA samples were thawed on ice. The cDNA was synthesized using the miRCURY LNA RT Kit (Qiagen) according to the manufacturer’s instructions, using 2 µL of RNA template per 10 µL total reaction volume. For cDNA synthesis quality control, 0.5 µL of a mixture of spike ins UniSp6 and cel-miR-39-3p (Qiagen) was added to the cDNA reaction. The cDNA samples were stored at 4°C when real-time quantitative polymerase chain reaction (RT-qPCR) was performed within four days after cDNA synthesis, or at −20°C when the RT-qPCR was performed at a later timepoint.

#### 2.1.3 RT-qPCR: Quality Control

For all samples, a quality control was performed to detect the presence of spike-ins UniSp2, UniSp4 and UniSp5 (to assess efficiency of miRNA isolation), of UniSp6 and cel-miR-39-3p (to assess efficiency of cDNA synthesis), and of the miRNAs miR-23a and miR-191 (to assess efficiency of endogenous miRNAs detection). To obtain miRNA-specific primers we checked the canine sequences on the miRBase database (16), and ordered miRCURY LNA miRNA PCR Assays (Qiagen; sequences and category numbers available through online data repository (17)). The cDNA samples were diluted 1:15, and 3 µL of diluted cDNA was used per well in a total reaction volume of 10 µL. Detection of targets was performed with miRCURY LNA SYBR® Green PCR Kits (Qiagen) according to the manufacturer’s instructions in a MyiQ^™^2 Two-Color Real-Time PCR Detection System (Bio-Rad, Veenendaal, The Netherlands).

#### 2.1.4 RT-qPCR: Custom plates

For the pilot study, miRCURY LNA miRNA Custom PCR Panels 96-well plates were ordered from Qiagen. These ready-to-use plates contained specific primer sets pre-coated in the wells. For each sample, 48 reactions were performed: UniSp3 (interplate calibrator for optimal calibration and control assessment), UniSp6 (cDNA spike-in control), one non-template control, five potential reference miRNAs, and 40 potential target miRNAs (Qiagen; sequences and category numbers available through online data repository (17)). Detection of targets was performed with miRCURY LNA SYBR® Green PCR Kits (Qiagen) according to the manufacturer’s instructions in a MyiQ^™^2 Two-Color Real-Time PCR Detection System (Bio-Rad). The threshold for detection of fluorescence was manually adjusted to 40 Relative Fluorescent Units to obtain the same threshold for each plate. The geometric mean of the five potential reference miRNAs was used to normalize the data. The relative expression of the target miRNAs was calculated using the 2^-ΔΔCT^ method (18).

### 2.2 Definitive study

#### 2.2.1 Animals

For the definitive study, we included blood samples of healthy dogs (*n* = 6), pre-operative blood samples of dogs with csACTs (*n* = 26), and both pre-and post-operative samples of dogs with PDH (*n* = 19). The blood samples from the pilot study were also included in the definitive study. Written informed consent was obtained from the owners for the participation of their animals in this study. For all dogs with PDH, surgery and sample collection were performed at the Faculty of Veterinary Medicine in Utrecht, as was the sample collection for the healthy dogs. For dogs with csACTs, the surgery and sample collection were performed at the Utrecht University Clinic or at external clinics. In case of external clinics, the samples were sent by post to the Faculty of Veterinary Medicine in Utrecht. For the healthy dogs both EDTA plasma and serum samples were available. For the dogs with PDH only EDTA plasma samples were available, while for the dogs with csACTs only serum samples were available. Because of these differences in sample types, miRNA expression levels were not compared between dogs with PDH and with csACTs, but only between dogs with either PDH or csACTs and healthy controls of corresponding sample types. Samples for the dogs with PDH and csACTs were collected between 2015 and 2020, samples for the healthy dogs were collected between 2019 and 2020. The clinical data of the dogs included in this study can be found in an online data repository (17).

Diagnoses of PDH and csACTs were made as described for the pilot study, as were the confirmations of healthy dogs. Recurrence of hypercortisolism after hypophysectomy was suspected based on recurrence of clinical signs, and confirmed by UCCR values above the reference range.

#### 2.2.2 RT-qPCR

Isolation of miRNAs from the blood samples and subsequent cDNA synthesis was performed as described for the pilot study. Quality control was first performed on all samples to determine efficiency of miRNA isolation, cDNA reaction, and detection of endogenously expressed miRNAs. For the definitive study, we pre-coated 384-well plates with a mixture containing the PCR primer mix (Qiagen; sequences and category numbers available through online data repository (17)), miRCURY LNA SYBR® Green PCR Kits (Qiagen), and nuclease-free water. The plates were stored at −20°C until further use. The assessed targets in the definitive study included five potential reference miRNAs and 15 potential target miRNAs, selected based on the pilot study. Detection of miRNAs was performed in a CFX384 Touch Real-Time PCR Detection system (Bio-Rad).

#### 2.2.3 Data analysis

To determine the pairwise variance and stability of miRNA expression, the geNorm (19) method from the SLqPCR package (v1.52.0, (20) using R (v3.6, (21) and RStudio (v1.3.1093, 21) was used. Data normalization was performed by subtracting the geometric mean of the CT values of the 12 most stably expressed miRNAs from the CT values of the target miRNAs (ΔCT). The relative expression of the target miRNAs was calculated using the 2^-ΔΔCT^ method (18). Normal distribution was assessed with the Shapiro-Wilk test. Because the data were not normally distributed, the Mann-Whitney U-test was performed to determine the significance of differences in expression between groups for independent samples, while paired sample analysis was performed with the Wilcoxon signed-rank test. The Spearman’s rank correlation coefficient test was used to assess correlation between variables. *P*-values <0.05 were considered significant. All statistical analyses were performed with SPSS Statistics for Windows (Version 27.0, IBM Corp, Armonk, NY, USA).

The datasets generated for this study can be found in the DataverseNL data repository (17, *made publicly available upon publication*).

## 3 Results

### 3.1 Pilot study

#### 3.1.1 Quality control

The quality control results (available through online data repository (17)) indicated that highly (mimicked by UniSp2), moderately (UniSp4), and lowly (UniSp5) expressed miRNAs could all be efficiently isolated and detected with our protocol, and that highly (UniSp6) and moderately (cel-miR-39-3p) expressed miRNAs were efficiently transcribed to cDNA. In addition, the detection of miR-23 and miR-191 in all the samples indicated that endogenously present miRNAs were also efficiently detected.

#### 3.1.2 Target miRNAs

Of the 40 target miRNAs assessed in the pilot study (results available through online data repository (17)), 6 were not detected in any samples (miR-34a-5p, miR-96-5p, miR-144-3p, miR-210-3p, miR-300-5p and miR-381-3p), while another 8 were not detected in most samples (miR-139-3p, miR-183-5p, miR-433-3p, miR-499a-5p, miR-455-5p, miR-483-5p, miR-497-5p, miR-499-5p). From the remaining 26 miRNAs, 15 miRNAs were selected for further analyses. The selected miRNAs showed potential differences in expression between patient groups or were potentially associated with the P/B value in the PDH group or with histopathological assessment in the csACT group.

### 3.2 Definitive study

#### 3.2.1 Stably expressed miRNAs

Five miRNAs were initially included in this study as normalization controls because we expected their expression levels to be stable: miR-23-3p, miR-191-5p, miR-222-3p, miR-423-5p and miR-425-5p. The selection of these miRNAs was based on endogenous miRNAs that are typically detected and stably expressed in human serum/plasma samples according to the Qiagen Guidelines for Profiling Biofluid miRNAs ©2019. We used geNorm software (19) to determine whether these miRNAs were indeed stably expressed in EDTA plasma samples (*n* = 44; 6 healthy dogs and 19 dogs with PDH, pre-op and post-op), in serum samples (*n* = 32; 6 healthy dogs and 26 dogs with csACT), and in the combined EDTA plasma/serum samples. The combination of miRNAs was interpreted as suitably stable when the Pairwise Variance (V-score) was lower than 0.15 (V_0.15_) (19,23). However, although the V-score decreased when all 5 miRNAs were used compared to when 3 or 4 miRNAs were used, they did not reach the V_0.15_ (Figure 1A).

**Figure 1.**
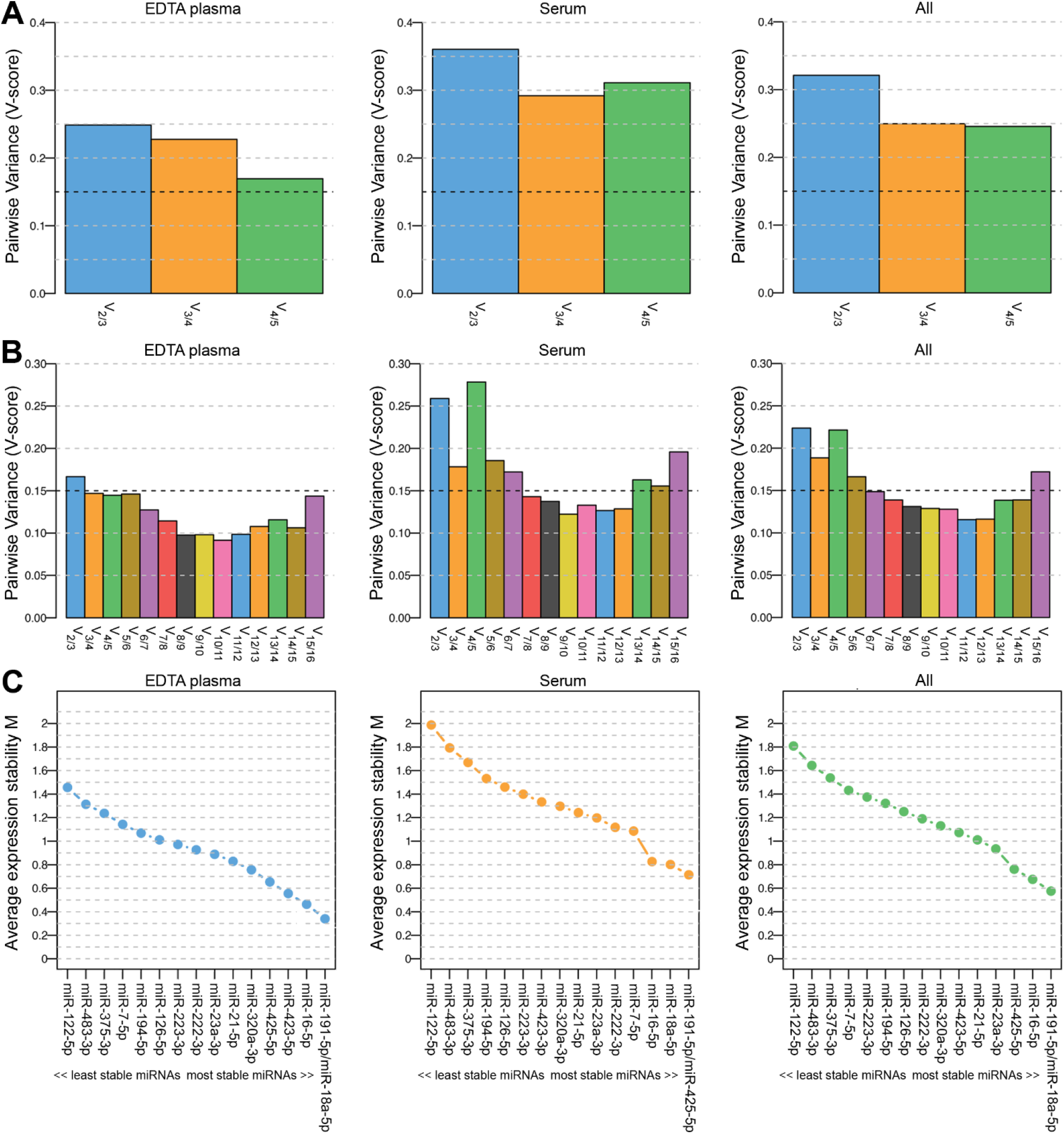
Pairwise Variance (V-score; A and B) and Average expression stability M (C) of miRNAs included in geNorm analyses to determine the optimal number of miRNAs for data normalization. Results when including 5 (A; miR-23a-3p, miR-191-5p, miR-222-3p, miR-423-5p and miR-425-5p) and when including 16 (B and C; miR-7-5p, miR-16-5p, miR-18a-5p, miR-21-5p, miR-23a-3p, miR-122-5p, miR-126-5p, miR-191-5p, miR-194-5p, miR-222-3p, miR-223-3p, miR-320a-3p, miR-375-3p, miR-423-5p, miR-425-5p and miR-483-3p) miRNAs. Left panels (EDTA plasma) include samples from healthy dogs (*n* = 6) and from dogs with pituitary-dependent hypercortisolism (*n* = 19, pre-op and post-op samples); middle panels (serum) include samples from healthy dogs (*n* = 6) and from dogs with cortisol-secreting adrenocortical tumors (*n* = 26); right panels (all) include both EDTA plasma and serum samples (*n* = 76).

To determine whether the V_0.15_ would be reached when more miRNAs were included in the analysis, we assessed the stability of all 20 assessed miRNAs, including the 15 intended target miRNAs. Only the miRNAs that could be detected in all samples were included, which left 16 miRNAs for the geNorm analysis (5 of the originally intended reference miRNAs and 11 target miRNAs). The lowest V-score (V_min;_ 0.115) in the combined EDTA plasma/serum samples group (“All”) was achieved with a combination of 12 miRNAs (Figure 1B). The most stably expressed miRNAs in both EDTA plasma and serum sample groups was miR-191-5p, while the least stably expressed miRNAs in all sample groups were miR-122-5p, miR-483-3p, and miR-375-3p (Figure 1C). Overall, both the V-scores (Figure 1B) and the Average expression stability M (Figure 1C) were lower in the EDTA plasma samples group than in the serum samples group.

For the subsequent analyses, we analyzed the expression levels of all 20 miRNAs. The geometric mean of the V_min_ (12 most stably expressed miRNAs) was used as normalization control.

#### 3.2.2 MiRNAs in Pituitary-Dependent Hypercortisolism

In the EDTA plasma samples, five miRNAs were expressed significantly higher in dogs with PDH (*n* = 19) compared to healthy dogs (*n* = 6): miR-122-5p (*p* = 2.9×10^−4^; no overlap between groups), miR-141-3p (*p* = 0.028), miR-222-3p (*p* = 0.008), miR-375-3p (*p* = 0.001), and miR-483-3p (*p* = 0.009) (Figure 2A). One miRNA was expressed significantly lower in dogs with PDH compared to healthy dogs: miR-126-5p (*p* = 0.036) (Figure 2A).

**Figure 2.**
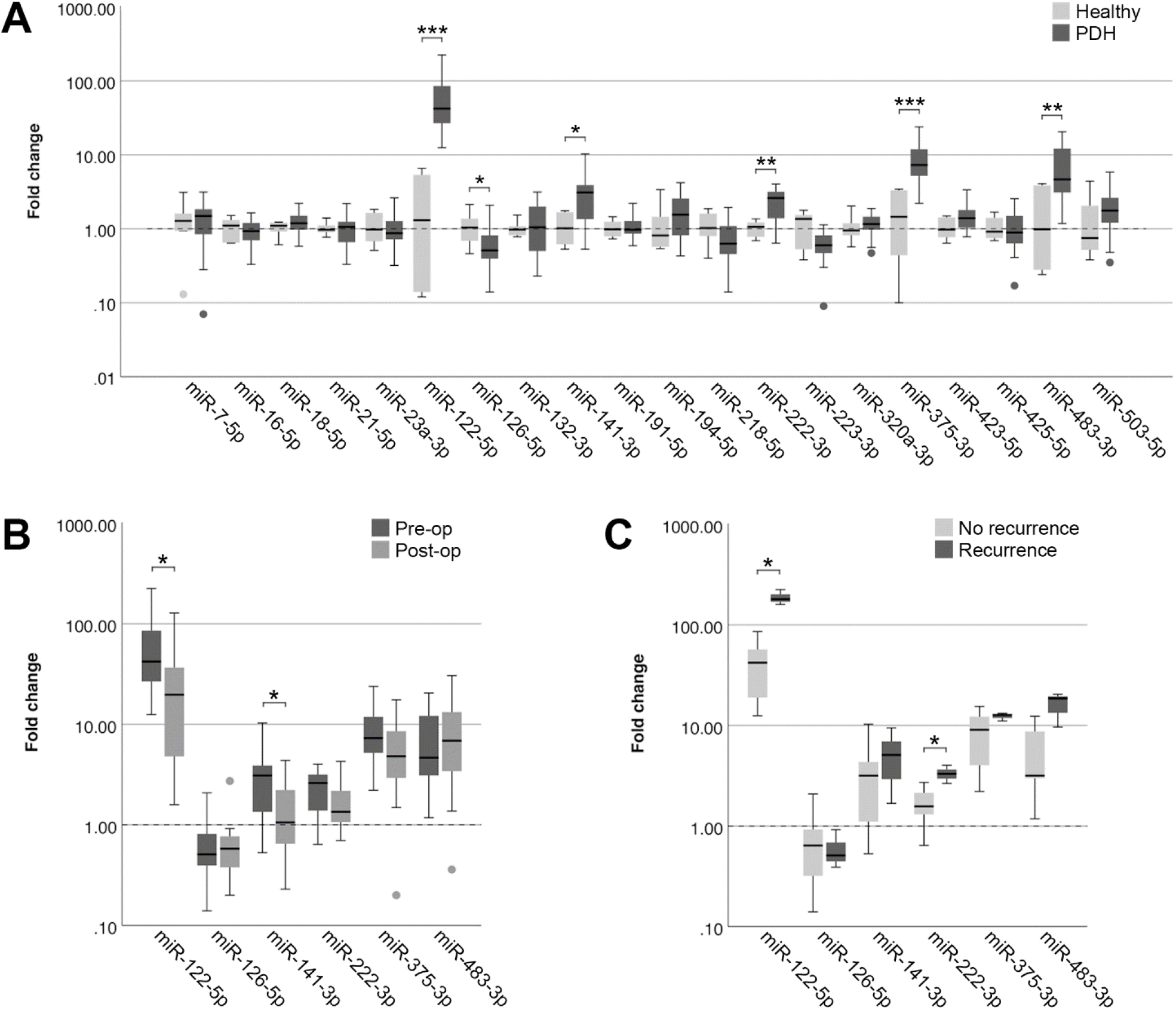
Circulating miRNA expression levels in EDTA plasma samples, normalized with V_min_ (12 most stably expressed miRNAs), in (A) dogs with PDH (*n* = 19) and healthy dogs (*n* = 6); (B) dogs with PDH (*n* = 19) before (pre-op) and after (post-op) hypophysectomy; and (C) dogs with PDH with (*n* = 3) or without (*n* = 7) recurrence of hypercortisolism within a year after hypophysectomy. Circles above and below boxes indicate outliers. * *p* < 0.05; ** *p* < 0.01; *** *p* < 0.001. PDH = pituitary-dependent hypercortisolism.

Of all 19 dogs with PDH, post-op samples were available. The timepoint of post-op sample collection was at median 3 days after hypophysectomy (range 1-7 days). The six differentially expressed miRNAs were analyzed in the post-op EDTA plasma samples, to determine whether their expression levels either decreased or increased after surgery. Paired-sample analyses of these six miRNAs showed that the expression levels of miR-122-5p (*p* = 0.013) and of miR-141-3p (*p* = 0.035) significantly decreased after surgery (Figure 2B). Although miR-222-3p and miR-375-3p showed a tendency to decrease based on Figure 2B, these changes were not significant (*p* = 0.117 and *p* = 0.165, respectively). The expression levels of miR-126-5p and miR-483-3p did not change in the post-op samples (*p* = 0.647 and *p* = 0.520, respectively).

##### 3.2.2.1 Association of miRNAs with recurrence of PDH

Because the P/B value is currently the most important predictor of recurrence after hypophysectomy, we analyzed whether expression levels of miRNAs (pre-operative samples) were correlated with the P/B value. None of the six differentially expressed miRNAs were significantly correlated with the P/B value, nor were any of the other fourteen miRNAs (Table 1).

**Table 1.** Correlation of pre-operative circulating miRNA expression with pituitary height to brain area value in dogs with PDH. Data analyzed with the Spearman’s rank correlation coefficient test.

|  | Correlation<br>coefficient (r) | Significance<br>(2-tailed; *p*) | Number (*n*) |
| --- | --- | --- | --- |
| miR-7-5p | 0.207 | 0.395 | 19 |
| miR-16-5p | 0.044 | 0.857 | 19 |
| miR-18a-5p | -0.029 | 0.908 | 19 |
| miR-21-5p | -0.083 | 0.737 | 19 |
| miR-23a-3p | -0.201 | 0.409 | 19 |
| miR-122-5p | 0.179 | 0.463 | 19 |
| miR-126-5p | -0.309 | 0.198 | 19 |
| miR-132-3p | -0.151 | 0.563 | 17 |
| miR-141-3p | 0.145 | 0.554 | 19 |
| miR-191-5p | 0.051 | 0.836 | 19 |
| miR-194-5p | 0.193 | 0.428 | 19 |
| miR-218-5p | -0.071 | 0.772 | 19 |
| miR-222-3p | 0.175 | 0.474 | 19 |
| miR-223-3p | 0.035 | 0.887 | 19 |
| miR-320a-3p | 0.247 | 0.307 | 19 |
| miR-375-3p | 0.067 | 0.786 | 19 |
| miR-423-5p | -0.116 | 0.637 | 19 |
| miR-425-5p | 0.105 | 0.668 | 19 |
| miR-483-3p | 0.023 | 0.926 | 19 |
| miR-503-5p | 0.068 | 0.788 | 18 |

Of the 19 dogs with PDH, follow-up information was available of 14 dogs. Of these dogs, 3 had recurrence of hypercortisolism (5, 7 and 12 months after hypophysectomy), 7 dogs had no recurrence of hypercortisolism and had follow-up of at least one year after surgery (median follow-up time 21 months, range 16 – 50 months), and 4 dogs had no recurrence of hypercortisolism but follow-up times of less than one year after surgery (median 5 months, range 2 – 8 months). The 3 dogs with reported recurrence were subsequently included in the “Recurrence” group, while the 7 dogs without recurrence and with follow-up time of at least one year after surgery were included in the “No recurrence” group. In assessing the expression profiles of the six differentially expressed miRNAs in the pre-operative samples, we found that miR-122-5p (*p* = 0.017) and miR-222-3p (*p* = 0.030) were expressed significantly higher in the dogs with recurrence than in the dogs without recurrence (Figure 2C).

##### 3.2.2.2 Differentially expressed miRNAs with one reference miRNA

Because the clinical application of miRNAs is limited when 12 miRNAs need to be assessed for data normalization, we wanted to determine whether the detected differences could still be observed when only one reference miRNA was used. As the most stably expressed miRNA in all sample groups, we chose to use the expression of miR-191-5p for data normalization. Although miR-126-5p lost its significance (*p* = 0.105), the other five miRNAs were still significantly higher in dogs with PDH compared to healthy dogs (Figure 3A) when normalized to only miR-191-5p. MiR-122-5p was equally significant when only one reference miRNA was used (*p* = 2.9×10^−4^) because still no overlap between sample groups was observed, but the *p*-values of the other four miRNAs increased (miR-141-3p: *p* = 0.049 vs 0.028; miR-222-3p: *p* = 0.026 vs 0.008; miR-375-3p: *p* = 0.003 vs 0.001; miR-483-3p: *p* = 0.022 vs 0.009).

**Figure 3.**
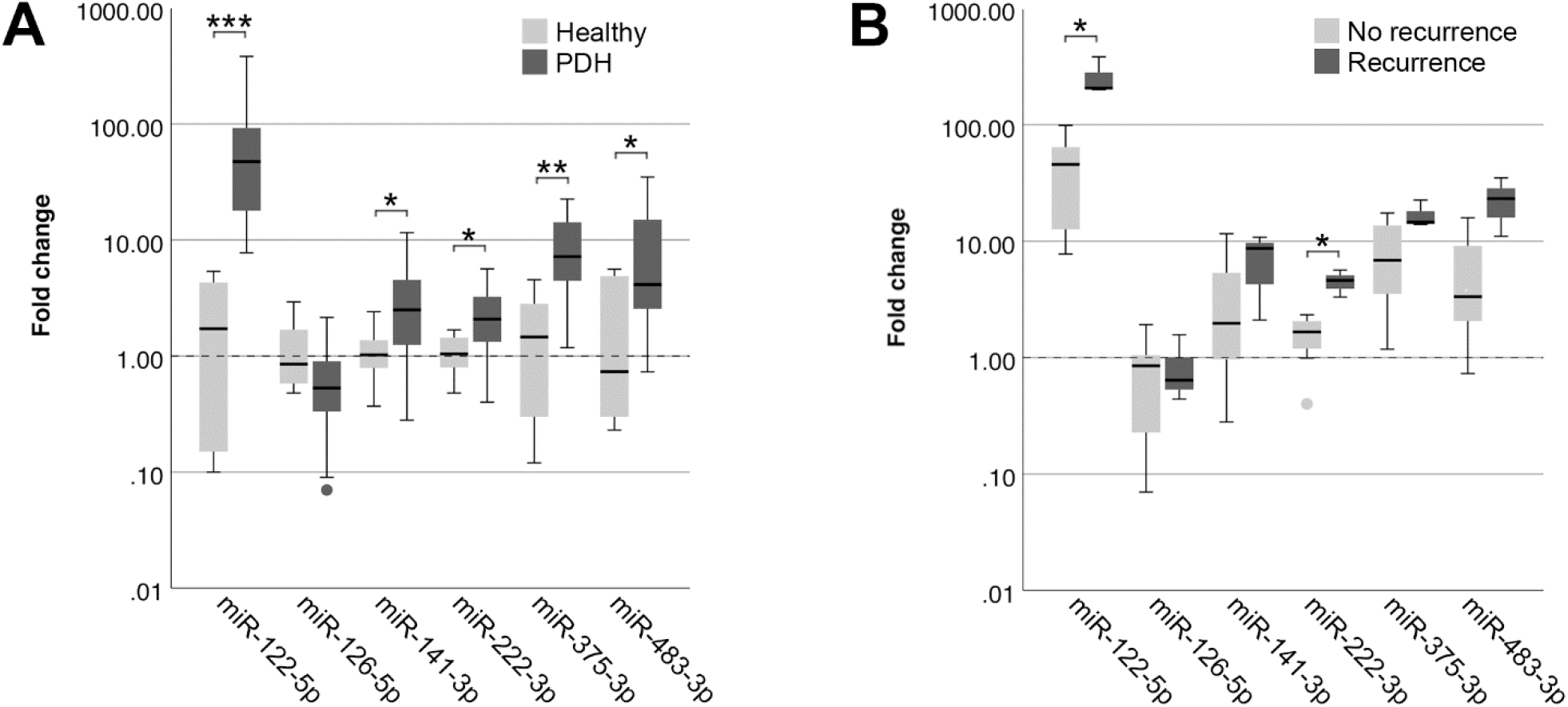
Circulating miRNA expression levels in EDTA plasma samples, normalized with only miR-191-5p, in (A) dogs with PDH (*n* = 19) and healthy dogs (*n* = 6); and (B) dogs with PDH with (*n* = 3) or without (*n* = 7) recurrence of hypercortisolism within a year after hypophysectomy. Circles below boxes indicate outliers. * *p* < 0.05; ** *p* < 0.01; *** *p* < 0.001. PDH = pituitary-dependent hypercortisolism.

When comparing the expression of these miRNAs between the recurrence group and the no recurrence group, normalized with only miR-191-5p, both miR-122-5p (*p* = 0.017) and miR-222-3p (*p* = 0.017) were still significantly higher in the recurrence group (Figure 3B).

#### 3.2.3 MiRNAs in Adrenal-Dependent Hypercortisolism

In serum samples, one miRNA was expressed significantly higher in dogs with a csACT (*n* = 26) compared to healthy dogs (*n* = 6): miR-483-3p (*p* = 0.020), while one miRNA was expressed significantly lower in dogs with a csACT: miR-223-3p (*p* = 0.038) (Figure 4A).

**Figure 4.**
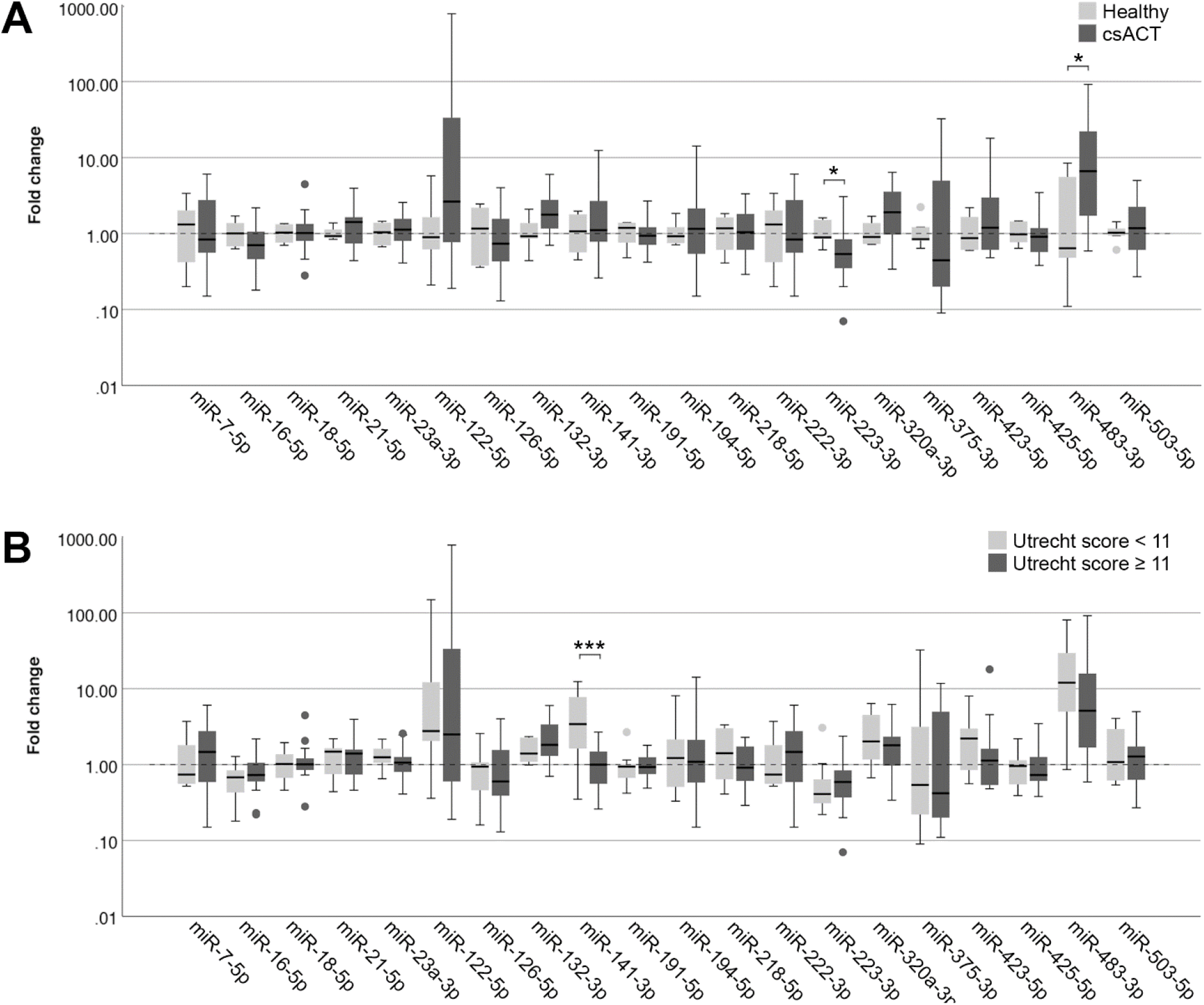
Circulating miRNA expression levels in serum samples, normalized with V_min_ (12 most stably expressed miRNAs), in (A) dogs with a csACT (*n* = 26) and healthy dogs (*n* = 6); and (B) dogs with a csACT with an Utrecht score below 11 (*n* = 9) compared to those with a score of 11 or more (*n* = 17). Circles above and below boxes indicate outliers. * *p* < 0.05; *** *p* < 0.001. csACT = cortisol-secreting adrenocortical tumor.

##### 3.2.3.1 Histopathological assessment

The median Ki67 PI in the csACTs (*n* = 26) was 8.2 % (range 0.3 – 50.7 %), and the median Utrecht score was 12.6 (range 4.3 – 57.7). Although miR-132-3p and miR-141-3p showed a moderate correlation with the Utrecht score, either positively (miR-132-3p) or negatively (miR-141-3p), these correlations were not significant (Table 2). None of the other miRNAs correlated to either the Ki67 PI or Utrecht score (Table 2).

**Table 2.**
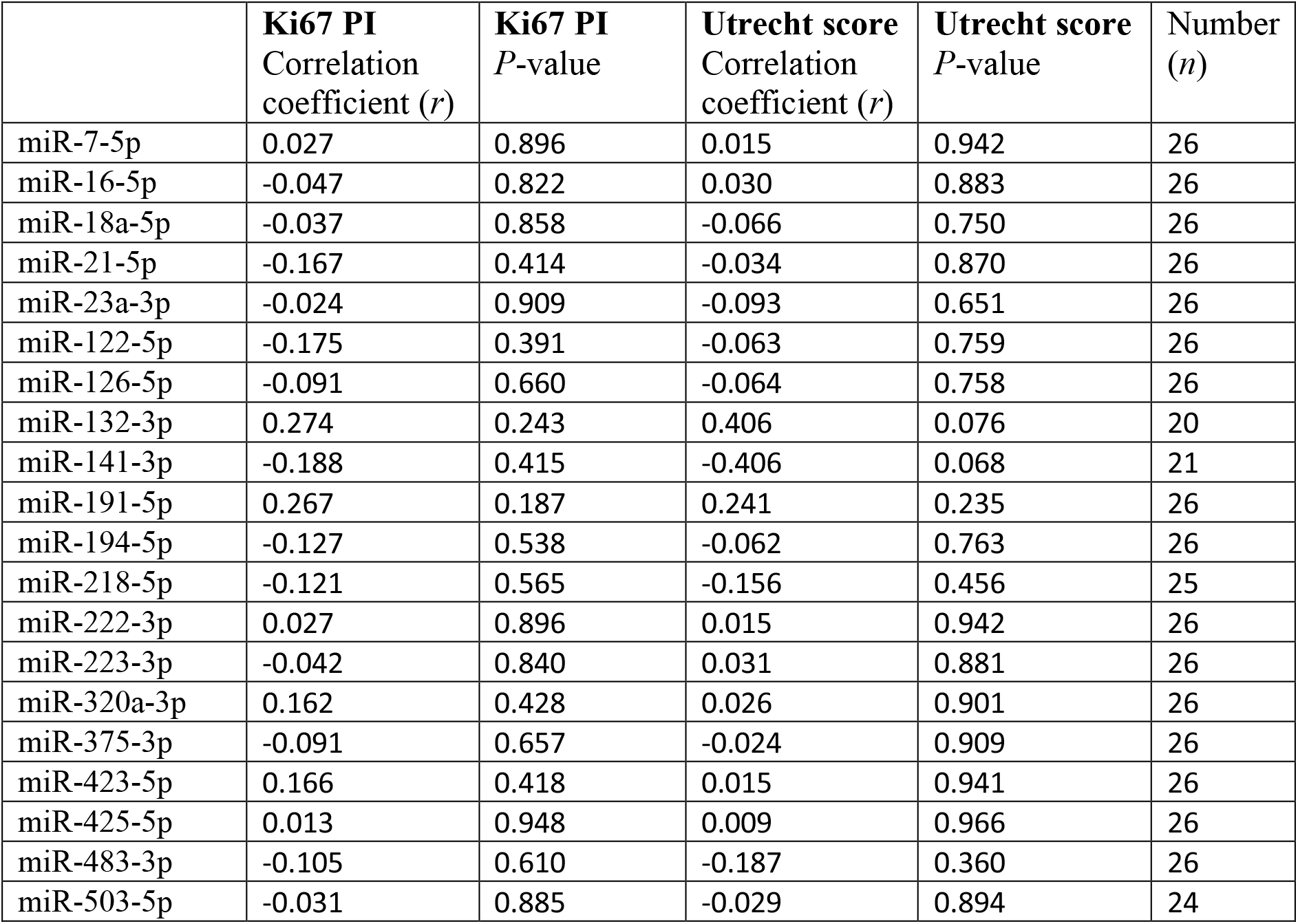
Correlation of circulating miRNA expression with Ki67 proliferation index (PI) and Utrecht score in dogs with csACTs. Data analyzed with the Spearman’s rank correlation coefficient test.

|  | **Ki67 PI**<br>Correlation<br>coefficient (*r*) | **Ki67 PI**<br>*P*-value | **Utrecht score**<br>Correlation<br>coefficient (*r*) | **Utrecht score**<br>*P*-value | Number<br>(*n*) |
| --- | --- | --- | --- | --- | --- |
| miR-7-5p | 0.027 | 0.896 | 0.015 | 0.942 | 26 |
| miR-16-5p | -0.047 | 0.822 | 0.030 | 0.883 | 26 |
| miR-18a-5p | -0.037 | 0.858 | -0.066 | 0.750 | 26 |
| miR-21-5p | -0.167 | 0.414 | -0.034 | 0.870 | 26 |
| miR-23a-3p | -0.024 | 0.909 | -0.093 | 0.651 | 26 |
| miR-122-5p | -0.175 | 0.391 | -0.063 | 0.759 | 26 |
| miR-126-5p | -0.091 | 0.660 | -0.064 | 0.758 | 26 |
| miR-132-3p | 0.274 | 0.243 | 0.406 | 0.076 | 20 |
| miR-141-3p | -0.188 | 0.415 | -0.406 | 0.068 | 21 |
| miR-191-5p | 0.267 | 0.187 | 0.241 | 0.235 | 26 |
| miR-194-5p | -0.127 | 0.538 | -0.062 | 0.763 | 26 |
| miR-218-5p | -0.121 | 0.565 | -0.156 | 0.456 | 25 |
| miR-222-3p | 0.027 | 0.896 | 0.015 | 0.942 | 26 |
| miR-223-3p | -0.042 | 0.840 | 0.031 | 0.881 | 26 |
| miR-320a-3p | 0.162 | 0.428 | 0.026 | 0.901 | 26 |
| miR-375-3p | -0.091 | 0.657 | -0.024 | 0.909 | 26 |
| miR-423-5p | 0.166 | 0.418 | 0.015 | 0.941 | 26 |
| miR-425-5p | 0.013 | 0.948 | 0.009 | 0.966 | 26 |
| miR-483-3p | -0.105 | 0.610 | -0.187 | 0.360 | 26 |
| miR-503-5p | -0.031 | 0.885 | -0.029 | 0.894 | 24 |

In our previously published introduction of the Utrecht score, we identified three groups with significantly different survival times after surgery when using cut-off values for the Utrecht score of < 6, ≥6 - < 11, and ≥ 11. In the current study, only one csACT had an Utrecht score of < 6, so for subsequent analyses we classified the csACTs as having an Utrecht score of < 11 (*n* = 9) or ≥ 11 (*n* = 17). When comparing miRNA expression between these groups, miR-141-3p was significantly higher in in the group with an Utrecht score of < 11 than in the group with an Utrecht score of ≥ 11 (*p* = .009, Figure 4B). None of the other miRNAs showed significant differences in expression between these groups (Figure 4B).

## 4 Discussion

Circulating miRNAs have been shown to be useful diagnostic and prognostic biomarkers in several diseases and cancer types (13,24). In this study, we identified several circulating miRNAs that are differently expressed in dogs with Cushing’s syndrome. The most clearly altered miRNA was miR-122-5p, which was significantly overexpressed in dogs with PDH and did not overlap with expression in healthy dogs. In addition, miR-122-5p expression was higher in dogs with recurrence after hypophysectomy than in dogs without reported recurrence. MiR-122-5p might therefore represent a useful noninvasive biomarker for dogs with PDH.

MiR-122^i^ is also of interest in other diseases and cancer types, both in humans and in dogs (25–29). In human cancers, the function of miR-122 seems to be dependent on its cellular context: while it has been described as an oncomiR in some cancers, including clear cell renal carcinoma (25) and colon cancer (29), it has been described as a tumor suppressor miRNA in others, including hepatocellular carcinoma (30) and bladder cancer (31). In several studies, miR-122 was found to inhibit the expression of aldolase, fructose-bisphosphate A (*ALDOA*) (29,32–34). ALDOA is a glycolytic enzyme that catalyzes the reversible conversion of fructose-1,6-bisphosphate to glyceraldehyde 3-phosphate and dihydroxyacetone phosphate. Although *ALDOA* is an important oncogene in several cancer types, including non-small cell lung cancer (35), pancreatic cancer (36) and hepatocellular carcinoma (37), it is a tumor suppressor in others, including prostate adenocarcinoma and stomach adenocarcinoma (37). The tumorigenic activity of ALDOA therefore seems to be context-dependent (37), and could potentially explain the context-dependency of miR-122. The expression of ALDOA has not yet been studied in pituitary tumors, which would be an interesting avenue for future research.

Although miR-122 expression has a clear link with cancer, there are also other diseases in which miR-122 expression can be dysregulated. Because miR-122 is derived from hepatocytes, it was found to be overexpressed in serum of dogs with different types of liver diseases (27,28,38). Because hypercortisolism can induce vacuolar hepatopathy (39), the increase in miR-122 levels seen in dogs with PDH could also be related to liver damage. In future studies, it would be interesting to determine whether miR-122 levels in dogs with PDH are correlated to their ALT values, as they were in a previous study on dogs with liver diseases (38). The finding that miR-122-5p levels in dogs with PDH significantly decreased in post-op samples compared to pre-op samples, even though the time between surgery and post-op sampling was relatively short (median 3 days, range 1-7 days) and the dogs still receive high doses of glucocorticoids for the first couple of days after hypophysectomy, suggests that its expression in dogs with PDH is not related to hypercortisolism but rather to the pituitary tumorigenesis. In addition, miR-122-5p levels were not significantly higher in dogs with hypercortisolism caused by a csACT than in healthy dogs. To validate the hypothesis that miR-122-5p expression is not hypercortisolism-related, it would be interesting to compare the miRNA expression levels that were found to be altered in this study in dogs with PDH to those of dogs with iatrogenic Cushing’s syndrome.

Another miRNA that was overexpressed in dogs with PDH compared to healthy dogs, as well as in dogs with PDH that had recurrence after hypophysectomy compared to dogs that did not have recurrence, is miR-222-3p. MiR-222 is overexpressed in many different human cancer types (40). A recent study that performed meta-analyses on 17 published articles on miR-222 showed that high expression of miR-222 significantly predicts poor overall survival in several human cancer types (41). MiR-222 could therefore also be a useful noninvasive biomarker in dogs with PDH.

Of the other four significantly altered miRNAs in PDH found in this study, only one was expressed significantly lower compared to healthy controls: miR-126-5p. MiR-126 has also been reported to be downregulated in other diseases in humans, including in diabetes mellitus (42). In contrast, circulating miR-126 was overexpressed in dogs with different types of neoplasia (43). Possibly, the miR-126-5p downregulation in the current study is related to the resulting endocrine syndrome, and not to the pituitary tumor. MiR-141-3p, miR-375-3p miR-483-3p were overexpressed in dogs with PDH and are also deregulated in several cancer types (44–46). For miR-375-3p, the difference in expression could also be related to the hypothalamic-pituitary-adrenal axis feedback loop: expression of miR-375 has been shown to be inhibited by corticotropin-releasing factor (CRF) (47). Since the high ACTH levels in dogs with PDH will result in low CRF levels (48), this lack of inhibition could result in higher miR-375p levels.

In dogs with csACTs, miR-483-3p was significantly overexpressed compared to healthy dogs. In humans, miR-483-5p, the reverse miRNA from the miR-483 locus, is one of the most commonly reported circulating miRNA that is overexpressed in patients with ACC compared to those with ACA (49–51). Although miR-483-3p is less studied, it is also overexpressed in ACC compared to ACA in humans (45). In the miRBase database, only the sequence for canine miR-483-3p is reported. Because of its importance in human ACTs, we nonetheless tested the primers for human miR-483-5p in our pilot study, but these did not result in a detectable product in the canine samples (17). Although the overexpression of miR-483-3p in dogs with csACTs compared to healthy dogs seems to be in line with its overexpression in human ACCs compared to ACAs, we did not see any differences in its expression when comparing csACTs with an Utrecht score of ≥ 11 with those that had an Utrecht score of < 11, nor was there a correlation of miR-483-3p expression with Ki67 PI or Utrecht score. Whether this is related to a difference in classification (e.g., the fact that we only included cortisol-secreting ACTs, or a potential overrepresentation of carcinomas in the total csACT group) or a difference in mechanism is currently unknown.

MiR-141-3p was expressed significantly lower in dogs with a csACT that had an Utrecht score of ≥ 11 than in those with an Utrecht score of < 11. MiR-141 has been reported as a useful biomarker in several cancer types, but was shown to be upregulated in some cancer types, including small cell lung cancer (44), and downregulated in others, including renal cell carcinoma (52). Recently, miR-141 was shown to inhibit angiogenesis both *in vitro* and *in vivo* (53). Previous research by our group showed that canine ACCs are in a more proangiogenic state compared to ACAs (54), which might be related to the decreased expression of miR-141-3p in the csACTs with an Utrecht score of ≥ 11. In future studies it would be interesting to determine whether miR-141-3p can predict the risk of recurrence after adrenalectomy.

MiRNAs that can predict malignancy or risk of recurrence before surgery would greatly impact clinical decision making. In dogs with PDH, both miR-122-5p and miR-222-3p were significantly overexpressed in dogs with recurrence after hypophysectomy compared to those without and are therefore interesting candidates as noninvasive biomarkers to predict recurrence. However, because only 3 dogs were included in the “Recurrence” group and 7 in the “No recurrence” group, these results will have to be validated in larger sample sizes. Additionally, in future studies it would be interesting to take blood samples at different times after surgery to determine the expression levels of these miRNAs at time of remission, and whether they will increase at time of recurrence before changes on diagnostic imaging are detectable. If so, these miRNAs could be highly useful early markers of recurrence.

The biggest hurdle in circulating miRNA analyses is data normalization. The five miRNAs that we originally intended to use as reference miRNAs were not stable enough, so we determined whether the stability would improve when we added all analyzed miRNAs in the geNorm analysis. Indeed, when adding more miRNAs, we were able to reach sufficiently stable results for data normalization. However, although this improves the reliability of our results in the current study, using 12 miRNAs for data normalization limits the clinical application of detecting circulating miRNAs. We therefore tested whether the differentially expressed miRNAs in dogs with PDH would retain their significance when normalized against only one miRNA. Although the *p*-values of some miRNAs slightly increased, in general the same results were observed as when using 12 miRNAs for normalization. Using only miR-191-5p, one of the two most stably expressed miRNAs in this study, as reference miRNA for data normalization therefore seems feasible in dogs with PDH. However, miR-191-5p can also be dysregulated in different cancer types (55,56), which makes it risky to use only one miRNA for data normalization. Ideally, to identify miRNAs that are stably expressed in dogs with different diseases, large-scale experiments should be performed that analyze expression of all circulating miRNAs with next generation sequencing in groups of dogs with different types of diseases. This would also help to identify novel useful miRNAs.

Overall, the miRNAs seemed to be more stably expressed in the EDTA plasma samples than in the serum samples as shown in the geNorm analyses. A previous study showed that there can indeed be differences in stability in EDTA plasma compared to serum samples (57). However, in the current study, handling of samples after collection could also have influenced the results. For example, all EDTA plasma samples from dogs with PDH were collected in our University Clinic and could be processed soon after collection, while samples from the dogs with csACTs were in several cases sent in from other clinics, so it took longer before the samples were processed and frozen down. Because of the small differences in EDTA plasma compared to serum samples, we decided not to directly compare the dogs with PDH to dogs with csACTs. For future studies it would be interesting to collect blood of all dogs in the same sample types, so that direct comparisons can be made.

To conclude, we have identified several circulating miRNAs that are dysregulated in dogs with Cushing’s syndrome. These miRNAs have the potential to be noninvasive biomarkers that may contribute to clinical decision-making.

## 5 Data availability statement

The datasets generated for this study will be made publicly available through the DataverseNL data repository (17).

## 6 Ethics statement

Ethical review and approval was not required for the study because no experimental interventions were carried out on the animals. Rest blood samples were collected when blood samples were taken for diagnostic purposes (dogs with PDH and csACTs) or preventive health screenings (healthy dogs). Written informed consent was obtained from the owners for the participation of their animals in this study.

## 7 Conflict of Interest

The authors declare that the research was conducted in the absence of any commercial or financial relationships that could be construed as a potential conflict of interest.

## 8 Author Contributions

KS and SG conceived and designed the experiments; KS, AV, AS and ET-S performed the experiments; KS, AV, HK, FR, BM and SG analyzed the data; HK, SD, FF, SvN and BM contributed materials; KS, AV and SG wrote the paper. All authors contributed to the article and approved the final version of the manuscript for submission.

## 9 Acknowledgements

We would like to thank Drs. Bart Sjollema, Rob Gerritsen, Giora van Straten, Jenny Buijtels and Dieneke Jongepier for contributing samples to our csACT dataset; Guy C.M. Grinwis for histopathological evaluations; all dog owners for letting us use their dogs’ samples for this study; Louis C. Penning for sharing his insights and expertise; and Friends of VetMed for their support.

## Footnotes

i When -3p or -5p miRNA was not specified in mentioned articles, only the miR-locus number will be mentioned

## Notes

https://doi.org/10.34894/XPIIU6

### Competing Interest Statement

The authors have declared no competing interest.

